# The ADAM17 sheddase complex regulator iTAP modulates inflammation, epithelial repair, and tumor growth

**DOI:** 10.1101/2022.04.11.487842

**Authors:** Marina Badenes, Emma Burbridge, Ioanna Oikonomidi, Abdulbasit Amin, Érika de Carvalho, Lindsay Kosack, Pedro Domingos, Pedro Faísca, Colin Adrain

**Affiliations:** Instituto Gulbenkian de Ciência (IGC), Oeiras, Portugal; Patrick G Johnston Centre for Cancer Research, Queen’s University, Belfast, UK; Genentech, South San Francisco, California, USA; Department of Physiology, Faculty of Basic Medical Sciences, University of Ilorin, Nigeria; Instituto de Tecnologia Química da Universidade Nova de Lisboa (ITQB-Nova), Oeiras, Portugal

**Keywords:** ADAM17/TACE, iRhom, iTAP/Frmd8, shedding, sheddase complex, inflammation, epithelium, tumor, epidermal growth factor receptor (EGFR), tumor necrosis factor (TNF).

## Abstract

The metalloprotease ADAM17 catalyzes the shedding of key signalling molecules from the cell surface, including the inflammatory cytokine TNF (tumour necrosis factor) and activating ligands of the EGFR (epidermal growth factor receptor). ADAM17 exists within an assemblage called the “sheddase complex” containing a rhomboid pseudoprotease (iRhom1 or iRhom2). iRhoms control multiple aspects of ADAM17 biology, including its vesicular trafficking, maturation from its precursor pro-form, activation on the cell surface and specificity for subsets of proteolytic targets. Previous studies from our laboratory and others identified the FERM domain-containing protein Frmd8/iTAP as an iRhom-binding protein. iTAP is required to maintain the cell surface stability of the sheddase complex, thereby preventing the precocious shunting of ADAM17 and iRhom2 to lysosomes and their consequent degradation. As pathophysiological role(s) of iTAP have not been addressed, here we sought to characterize the impact of loss of iTAP on ADAM17-associated phenotypes in mice. Our data show that iTAP KO mice exhibit defects in ADAM17 activity in inflammatory and intestinal epithelial barrier repair functions, but do not exhibit the collateral effects associated with global loss of ADAM17. Furthermore, we show that iTAP promotes cancer cell growth in a cell-autonomous manner, and by modulating the tumor microenvironment. Our work suggests that pharmacological intervention at the level of iTAP may be beneficial to target ADAM17 activity in specific compartments during chronic inflammatory diseases or cancer, avoiding the deleterious impact on vital functions associated with the widespread inhibition of ADAM17 in normal tissues.

## Introduction

The release of signalling proteins from the plasma membrane often involves the so-called “shedding”: the proteolytic cleavage of the extracellular domains of integral membrane proteins, catalysed by membrane-tethered metalloproteases (Lichtenthaler *et al*., 2018). This process of combining a membrane-anchored substrate and a membrane-tethered protease enables the strict spatiotemporal control over signalling that is required during development, homeostasis and immune responses. ADAM17 is a prominent sheddase that cleaves/releases a range of important signalling molecules including the apical inflammatory cytokine TNF (tumour necrosis factor), activating ligands of the EGFR (epidermal growth factor receptor) and key cell adhesion molecules including L-selectin (Zunke and Rose-John, 2017). ADAM17 substrates control multiple important biological processes including inflammation, growth control, development, metabolism and cell adhesion (Zunke and Rose-John, 2017).

In recognition of the importance of ADAM17 for inflammatory disease and cancer, there have been numerous attempts by pharma to develop safe ADAM17 inhibitors (Calligaris *et al*., 2021). However, these attempts have failed at (or before) phase II trials because of safety concerns caused by collateral targeting of other members of the wider metzincin metalloprotease subgroup that share a common active site architecture with ADAM17 (Murumkar *et al*., 2020). New approaches are therefore needed to identify strategies to safely block the cleavage of ADAM17 substrates during disease, ideally targeting a specific tissue or compartment.

Over the past 10 years, significant knowledge has emerged concerning how ADAM17 is regulated at the biochemical, cell biological and organismal levels. We and others discovered that certain pseudoenzymes called iRhoms are essential cofactors for ADAM17; without iRhoms ADAM17 is proteolytically inactive and hence ADAM17 substrate shedding is abolished (Adrain *et al*., 2012; McIlwain *et al*., 2012; Siggs *et al*., 2012; Li *et al*., 2015) (Adrain and Cavadas, 2020). Mammals encode two iRhom paralogs with partially redundant roles in ADAM17 regulation at the organismal and cellular levels (Christova *et al*., 2013; Li *et al*., 2015). iRhom1 regulates ADAM17 shedding in the brain(Sun *et al*., 2021), nervous system (Tüshaus *et al*., 2021) and in endothelial cells (Babendreyer *et al*., 2020). iRhom2 KO which develop normally but fail to secrete TNF, a key ADAM17 substrate that coordinates the responses to infection and chronic inflammatory diseases; loss of iRhom2 attenuates the development of multiple inflammatory disease models in mice (Adrain *et al*., 2012; McIlwain *et al*., 2012; Siggs *et al*., 2012; Adrain and Cavadas, 2020) (Barnette *et al*., 2018) (Kim *et al*., 2020) (Issuree *et al*., 2013; Luo *et al*., 2016; Chaohui *et al*., 2018; Qing *et al*., 2018; Sundaram *et al*., 2019). By contrast, the double KO of iRhom1 and iRhom2 in mice results in embryonic or perinatal lethality (Christova *et al*., 2013; Li *et al*., 2015).

This tissue-specific regulation of ADAM17 by different iRhom proteins has appealing therapeutic implications, since if it were possible to specifically target an individual iRhom (e.g. iRhom2), this would avoid the severe collateral effects associated with global inhibition of ADAM17 activity. Hence, targeting iRhom2 may enable specific inhibition of the inflammatory properties of ADAM17, while avoiding the global inhibition of its activity.

iRhom proteins physically interact with ADAM17, forming an assemblage called the “sheddase complex” (Künzel *et al*., 2018; Oikonomidi *et al*., 2018) and fulfil many key roles in ADAM17 regulation. This includes the vesicular transport of ADAM17 from its site of biogenesis, the endoplasmic reticulum (ER), into the *trans*-Golgi (Adrain *et al*., 2012; McIlwain *et al*., 2012) where ADAM17 undergoes a crucial activation step: the removal of its inhibitory prodomain by pro-protein convertases (Schlöndorff *et al*., 2000). Therefore, ADAM17 is catalytically inactive in iRhom KO cells because it fails to exit the ER and cannot undergo this critical maturation step (Adrain *et al*., 2012). Follow up studies from several groups later found that ADAM17 remains complexed to iRhom proteins on the cell surface, where iRhoms act as a signal-sensing scaffold within the sheddase complex (Cavadas *et al*., 2017; Grieve *et al*., 2017). Upon receipt of suitable signalling cues (e.g. phorbol esters, inflammatory stimuli), the cytoplasmic tail of iRhom proteins undergo phosphorylation, which drives the recruitment of 14-3- 3 proteins to this domain of iRhom (Cavadas *et al*., 2017; Grieve *et al*., 2017). This event triggers the activation of ADAM17, potentially driven by triggering a molecular rearrangement of the complex or displacing ADAM17 from the sheddase complex (Adrain and Cavadas, 2020).

To understand in more detail how the sheddase complex is regulated, we and others performed immunoprecipitation experiments coupled to mass-spectrometry to isolate iRhom-binding partners. This identified a largely uncharacterised protein that we named iTAP (iRhom-tail-associated protein), also called Frmd8 (Künzel *et al*., 2018; Oikonomidi *et al*., 2018). iTAP is a member of the FERM (Band 4.3 protein, Ezrin, Radixin, Moesin) superfamily, whose canonical function is anchoring membrane protein “clients” to cortical actin at the plasma membrane (Chishti *et al*., 1998; Fehon *et al*., 2010). Consistent with this general role for FERM domain proteins, iTAP binds to the cytoplasmic tail of iRhom proteins and it is essential to maintain the stability of the sheddase complex on the plasma membrane(Künzel *et al*., 2018; Oikonomidi *et al*., 2018). iTAP ablation triggers the selective loss of mature (active) ADAM17 in a range of iTAP KO cells and mouse tissues, because of precocious mis-sorting of the sheddase complex to lysosomes where it is degraded(Künzel *et al*., 2018; Oikonomidi *et al*., 2018).

Like iRhom2 KOs, iTAP KO mice are born at normal mendelian ratios, appear normal and are fertile (Künzel *et al*., 2018; Oikonomidi *et al*., 2018). In the present study we sought to characterize the impact of loss of iTAP on ADAM17-associated phenotypes in mice. Our data suggest that iTAP KO mice do not exhibit the collateral defects associated with global loss of ADAM17 but exhibit defects in inflammatory and intestinal epithelial barrier functions that have been reported in ADAM17 mutant models. Furthermore, we show that iTAP promotes cancer cell growth in a cell-autonomous manner and by modulating the tumor microenvironment. Our work identifies that intervention at the level of iTAP may be beneficial to target the inflammatory features of ADAM17 associated with iRhom2, and shows new insights of iTAP-mediated sheddase complex on cancer growth regulation.

## Results

### iTAP KO mice exhibit loss of mature ADAM17 in multiple tissues and reduced shedding of the ADAM17 substrate L-selectin

To understand better the role of iTAP as a regulator of the sheddase complex, we and others previously generated Frmd8/iTAP-null mice (Künzel *et al*., 2018; Oikonomidi *et al*., 2018). *Frmd8* homozygous mutant mice are born at normal mendelian ratios, are viable, fertile, and do not present any gross anatomical abnormalities compared to WT littermates (Künzel *et al*., 2018; Oikonomidi *et al*., 2018). However, as these previous studies did not exclude the possibility of subtle phenotypes, we carried out a detailed characterisation of iTAP KO mice. In some, but not all (Li *et al*., 2007; Gelling *et al*., 2008) ADAM17 KO mouse studies, ADAM17-null mice exhibit perinatal lethality and pronounced epithelial defects in multiple tissues, many of which are associated with defective EGFR signalling (Peschon *et al*., 1998), including precocious eyelid opening, defects in eye, lung, and heart valve morphogenesis, and deranged hair follicles with irregular pigment deposition (Peschon *et al*., 1998). Many of these phenotypes are also observed in iRhom1/iRhom2 double KO (DKO) embryos(Li *et al*., 2015), but, aside from defective inflammatory responses, iRhom2-null mice only present defects in keratinocyte proliferation, presenting thinner footpads and irregular pigment deposition (Maruthappu *et al*., 2017). We found that iTAP KO embryos develop normally, are born with their eyes closed as per WT littermates, and exhibit no obvious morphological differences in any of the major organs surveyed (Fig. S1A,B). Like iRhom2 KOs (Adrain *et al*., 2012), iTAP KOs lacked ADAM17-associated defects in the eye at E14.5 (Fig. S1B) while other epithelial tissues (small intestine, colon, caecum, lung) and the heart Fig. S1C,D) exhibited no obvious differences compared to controls.

Deletion of iTAP triggers the loss of the lower molecular weight form of ADAM17 that corresponds to the mature (i.e., active) form of the protease (Schlöndorff *et al*., 2000), as a consequence of its precocious trafficking to and degradation in lysosomes (Künzel *et al*., 2018; Oikonomidi *et al*., 2018). Interestingly, isolated primary keratinocytes from iTAP KOs exhibited substantially reduced levels of mature ADAM17 (Fig. S1E); however, some residual amounts of mature ADAM17 remained (Fig. S1E). These residual levels of mature ADAM17 may potentially explain the lack of overt histological skin or hair follicle density defects in iTAP KOs (Fig. S1F-H). Finally, as a hypermetabolic phenotype has been reported for the minority of ADAM17 KOs that survive perinatal lethality until adulthood(Gelling *et al*., 2008), we tested the growth rate and food intake of iTAP KO juveniles. While this does not preclude differences upon metabolic challenge, iTAP KO mice appeared normal under steady state conditions (Fig. S1I and S1J). In summary, like iRhom2 KO mice but unlike ADAM17 KO or iRhom1/iRhom2 DKO animals, iTAP KOs do not present overt growth or developmental defects.

As our previous studies established that ablation of iTAP in primary human macrophages results in a pronounced impairment of TNF shedding by ADAM17 (Oikonomidi *et al*., 2018), here we focused on the physiological impact of iTAP loss on the protein levels and sheddase activity of ADAM17 in immune cells of mice, in which ADAM17 and iRhom2 play a prominent role (Horiuchi *et al*., 2007; Li *et al*., 2007; Adrain *et al*., 2012; McIlwain *et al*., 2012; Siggs *et al*., 2012). Strikingly, we observed that iTAP-deficient splenocytes and lymphocytes exhibited substantially reduced levels of mature ADAM17 (Fig. 1A,B). We also observed substantial depletion of mature ADAM17 in B cells isolated from spleen or lymph nodes, as well as in dendritic cells differentiated *ex vivo* from bone marrow progenitors (Fig. S2A-C; gating strategy for B cell sorting indicated in Fig. S3A-E). To further probe the impact of iTAP depletion on ADAM17 activity in immune cells, we assessed the shedding of the endogenous ADAM17 substrate L-selectin (Marczynska *et al*., 2014) in response to the ADAM17 shedding stimulant, PMA. Notably, the PMA-triggered shedding of L-selectin was substantially impaired in T cells (Fig. 1C-E; gating strategy indicated in Fig. S4A-H) and to a more modest extent in B cells (Fig 1F; gating strategy indicated in Fig. S4A- D, I). Consistent with a role for defective ADAM17 activity, L-selectin shedding could be rescued by the broad-spectrum metalloprotease inhibitor Marimastat but was insensitive to the ADAM10-selective inhibitor GI 254023X (Fig. 1C-F). In summary, together with previous studies (Künzel *et al*., 2018; Oikonomidi *et al*., 2018), our data indicate that iTAP is a physiological regulator of mature ADAM17 levels and activity in immune cells.

**Figure 1.**
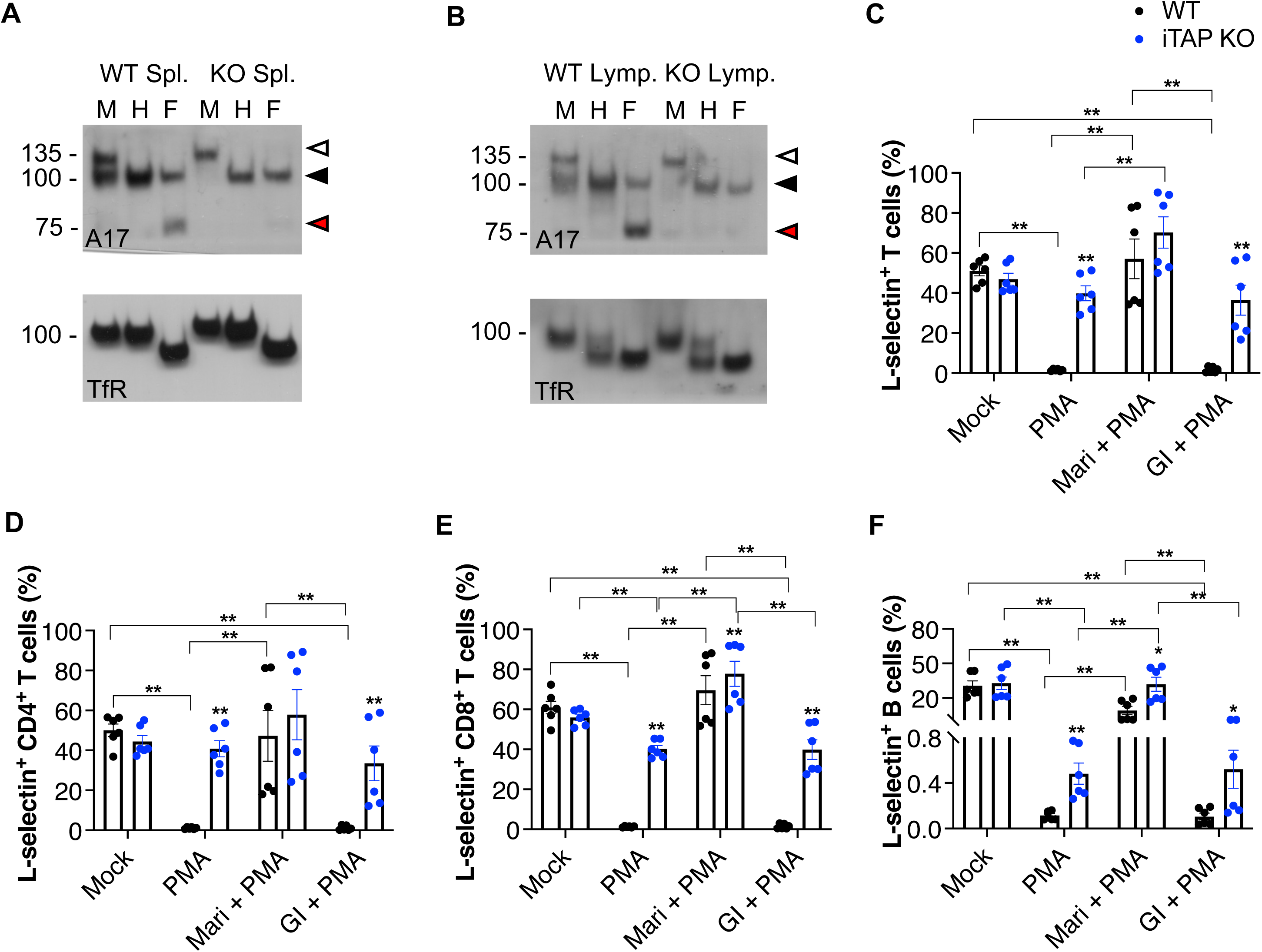
iTAP KO mice have defects in ADAM17 maturation and shedding. **A-B** Immunoblots showing immature versus mature ADAM17 (A17) in (A) splenocytes and (B) lymphocytes from WT versus iTAP KO mice. Protein samples were either mock-treated (M) or deglycosylated with Endo-H (H) or PNGase F (F). Immunoblots for the Transferrin receptor (TfR) serve as a loading control. n=2 The immature form of ADAM17 is indicated by white arrowheads; the black arrowhead denotes both glycosylated mature ADAM17 and deglycosylated immature ADAM17 respectively (which have similar electrophoretic mobility), whereas red arrowheads denote the fully deglycosylated, mature, ADAM17 polypeptide. **C-F** Levels of L-selectin in the surface of T cells (CD45.2^+^, CD3^+^), (C), CD4-positive T cells (CD45.2^+^, CD3^+^, CD4^+^) (D), CD8-positive T cells (CD45.2^+^, CD3^+^, CD8^+^) (E) B cells (CD45.2^+^, B220^+^) (F) isolated from the spleen of the mice described above. The isolated splenocytes were pre-treated with DMSO, ADAM17 inhibitor Marimastat (Mari) or ADAM10 inhibitor GI254023X (GI) and then treated with DMSO or PMA. n=2 with 3 replicates per experiment. Results are indicated as mean ± SEM. * represents p<0.05, ** represents p<0.01.

### iTAP KOs exhibit ADAM17 defects in experimental sepsis and colitis models

As noted above, iRhom2 plays an essential role in the trafficking regulation and sheddase activity of ADAM17 in immune cells, particularly macrophages–a key TNF source *in vivo*, the major inflammatory cytokine whose release requires the ADAM17/iRhom2 sheddase complex (Adrain *et al*., 2012; McIlwain *et al*., 2012; Siggs *et al*., 2012). To discern the requirement for iTAP for inflammatory cytokine release, we challenged iTAP KOs with two inflammatory models. We found that in response to LPS stimulation, bone marrow-derived macrophages differentiated *ex vivo* from iTAP KO mice exhibited substantially less TNF shedding than WT counterparts, while, as expected from iRhom KO studies (Adrain *et al*., 2012), there was no alteration in the levels of the soluble cytokine IL-6, whose release does not require ADAM17 (Fig 2A,B). To probe this observation in an *in vivo* context, we challenged iTAP KO mice in a sepsis model driven by *i.p*. administration of LPS (Fig. 2C,D). Similar to the *in vitro* studies, we found that iTAP KO mice have significantly less serum TNF, but as anticipated, unaltered IL-6 levels compared to controls (Fig. 2C,D). Hence, similar to iRhom2 KO or ADAM17 mutant mice, iTAP controls the shedding of TNF *in vivo*.

**Figure 2.**
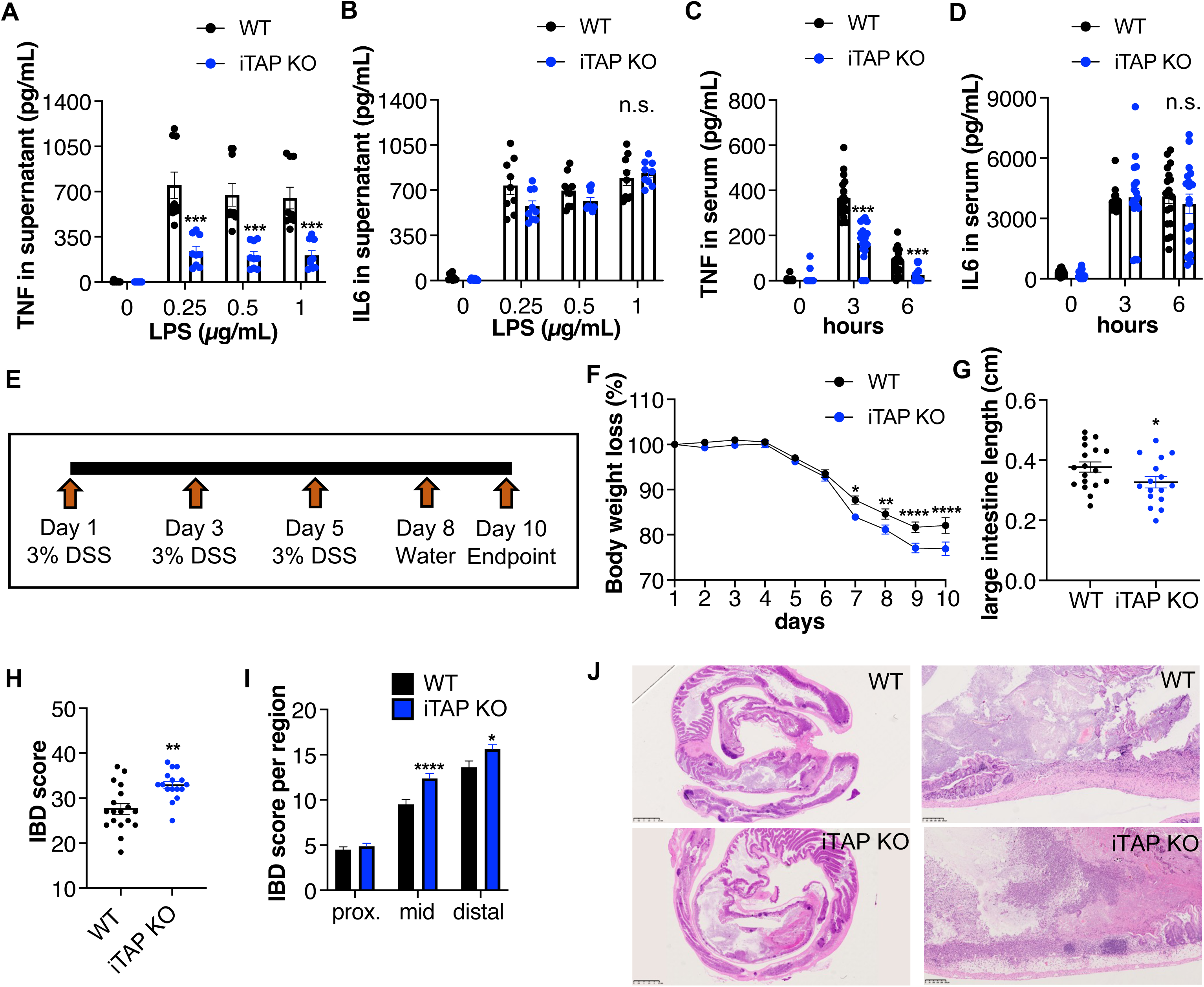
iTAP KO mice have ADAM17-associated defects in TNF shedding and epithelial regeneration. **A-B** Level of TNF (A) and IL-6 (B) secretion in culture supernatants from primary macrophages isolated from WT versus iTAP KO mice and stimulated with 3 doses of LPS (0.25, 0.5 and 1 μg/mL) for 2h (TNF) or 6h (IL-6). n=3 with 3 replicates per genotype per experiment. **C-D** TNF (C) and IL-6 (D) levels in the serum of WT versus iTAP KO mice following *i.p.* injection of PBS (0) or LPS (37,5 μg/g) for 3 or 6h. n= 3 with 6 mice per genotype per condition. **E** Schematics of the DSS-driven acute colitis model. **F-K** Body weight loss (%) (F), and large intestine length (cm) (G) of WT versus iTAP KO mice after DSS administration. **H-J** Histopathological IBD score in the total area (H) or by section (proximal, mid and distal) (I) and H&E images of the large intestine (J) of the mice described above. n= 3 with 6 mice per genotype per experiment. Results are indicated as mean ± SEM. Scale bar is indicated within the H&E images. * represents p<0.05, ** represents p<0.01, *** represents p<0.001, **** represents p<0.0001.

While ADAM17 and iRhom2 are required for the propagation of inflammatory responses, the predominant role for ADAM17 in the intestinal epithelium appears to be the promotion of epithelial repair (Chalaris *et al*., 2010). Hence, ADAM17 mutants are sensitized to intestinal damage driven by an experimental DSS (dextran sulfate sodium)-induced colitis model (Chalaris *et al*., 2010). Consistent with this ADAM17 phenotype, iTAP KO mice were more prone to worsened indicators of disease triggered by a model of acute DSS-induced colitis (Fig. 2E), including more pronounced body weight loss (Fig. 2F), and colon shortening (Fig 2G). Associated with this, iTAP KOs exhibited more severe disease in a histopathological-based score of disease severity (IBD score (Seamons *et al*., 2013)), particularly in the mid-colon, with more pronounced mucosal loss and more area of the intestine affected (Fig. 2H-J). Our data hence establish that like ADAM17, iTAP is required for intestinal regeneration/repair *in vivo*. Notably, evidence supporting a role for iRhom2 in this or similar paradigms is limited: whereas iRhom2 KO mice are not sensitized to DSS-induced colitis (Siggs *et al*., 2012; Geesala *et al*., 2019) they are nevertheless, in fact sensitized to a model of colitis driven by the loss of IL10 and exhibit spontaneous colitis in this context, which has been proposed to be associated with intestinal inflammation and microbiota-associated perturbations (Geesala *et al*., 2019).

### iTAP promotes tumour growth from within the tumor microenvironment

ADAM17 is crucial for the shedding of multiple substrates that can promote a state of chronic inflammation, contributing to the development and/or dissemination of cancer, such as TNF, or a soluble form of the IL6 receptor (Miller *et al*., 2017; Düsterhöft *et al*., 2019). Tumor cells often present increased TACE expression leading to autocrine EGFR activation (Borrell-Pagès *et al*., 2003) (Dong *et al*., 1999; Gschwind *et al*., 2003) (Schumacher and Rose-John, 2022). In addition, ADAM17 is crucial for TNF/TNFR1- mediated necroptosis, and for tumor cell induced endothelial cell death, which facilitates tumor cell extravasation and metastasis(Bolik *et al*., 2022). As ADAM17 maturation is decreased in multiple tissues of iTAP KO mice, including the lung (Oikonomidi *et al*., 2018), and defects in its sheddase activity were observed in *ex vivo* (Fig 1C-F), and in *in vivo* experiments (Fig. 2) we next evaluated the effect of iTAP depletion on tumour growth and metastasis. We used an allograft model based on the subcutaneous injection of Lewis lung carcinoma (LLC) tumor cells into immunocompetent WT *versus* iTAP KO mice (Fig. 3A), to assess the impact of iTAP loss in the tumor niche on the growth of WT LLC cells (Mendonça *et al*., 2019). Strikingly, we found that iTAP KO host mice were protected from tumor growth, resulting in tumors of a lower volume and mass (Fig. 3B-E). Consistent with a role for iTAP in the dissemination of inflammatory responses, we observed smaller necrotic lesions (Fig. 3F) and reduced superficial ulceration (data not shown) in tumors from iTAP KO hosts, as well as decreased levels of the mRNAs for inflammatory cytokines (TNF, IL6 and IL1β) and chemokines (CXCL2 and CCL4) compared to tumors from WT mice (Fig. 3G). In this model of subcutaneous-mediated LLC cell inoculation, we did not detect significant metastases to the lungs (a site where LLC cells typically accumulate in other models). However, consistent with the reduced subcutaneous topical growth of the tumors in iTAP KO hosts, we observed a trend towards a reduced number of micrometastases in iTAP KO lungs (Fig. 3H,I). Similarly, although inflammatory infiltrates were modest in the lungs of mice of both genotypes, we nevertheless found reduced TNF and IL1β mRNA levels in the lungs of the KOs compared to controls (Fig. 3J). Taken together, our data reveal that iTAP expression in the tumor microenvironment can promote local and metastatic tumor growth.

**Figure 3.**
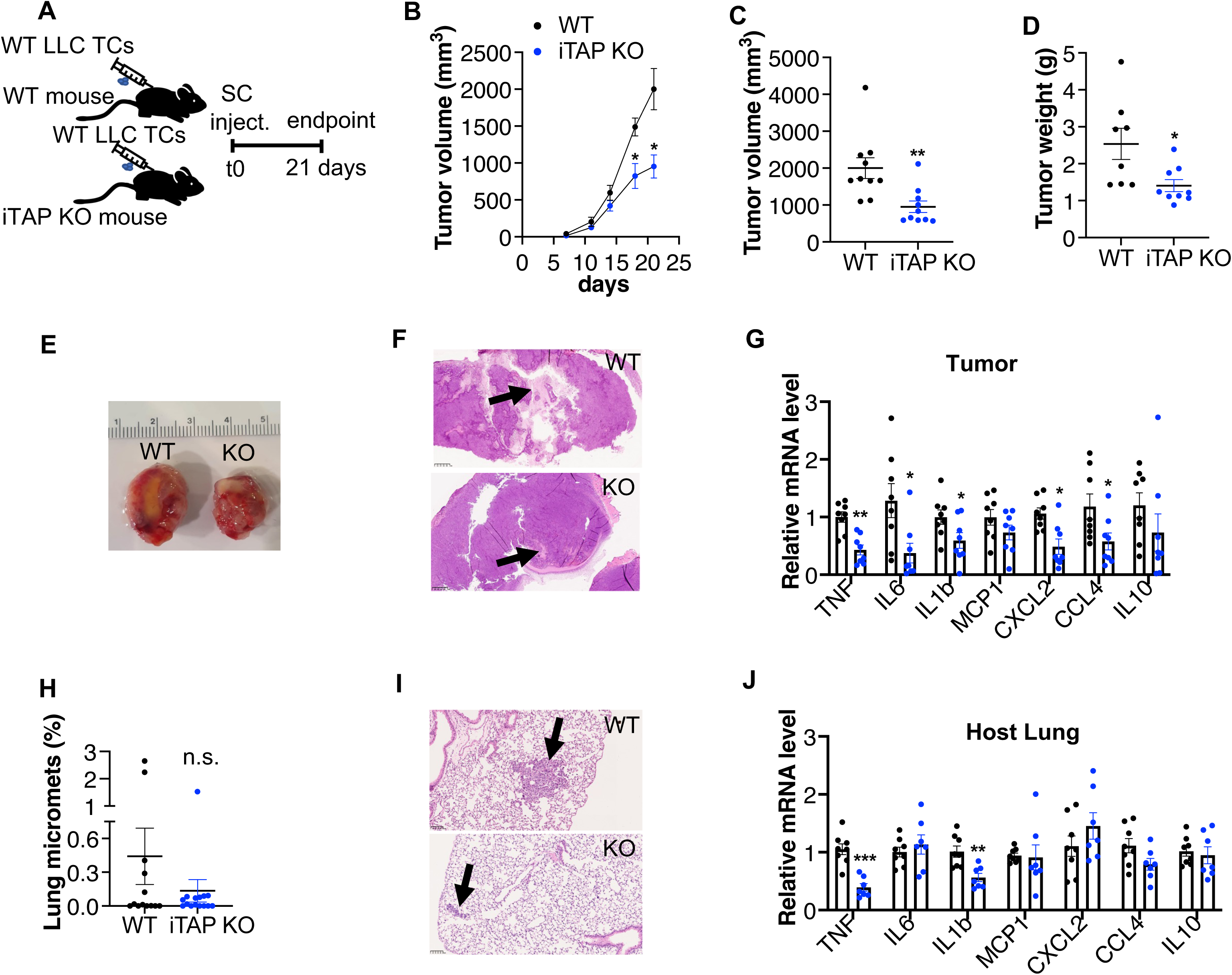
iTAP expression in the microenvironment promotes tumor growth. **A** Schematic illustrating the tumor model based on subcutaneous injection of WT LLC cells into WT versus iTAP KO immunocompetent hosts. **B-E** LLC cell-derived tumor volume over time (B) and at the endpoint (C), tumor weight (D), and representative photographs of the tumors (E) isolated from WT and iTAP KO mice. **F** Images of necrotic areas of the tumors described above. **G,J** Relative mRNA expression of inflammatory genes in the tumors (G) and lungs (J). Expression was normalized to the level of the housekeeping control *Gapdh*. **H, I** Percentage of micrometastases (H) and representative H&E-stained images (I) of the lungs of the animals described above. n=3 with 2-5 mice per genotype per experiment. Results are indicated as mean ± SEM. Scale bar is indicated within the H&E images. * represents p<0.05, ** represents p<0.01, *** represents p<0.001.

### Cell-autonomous expression of iTAP promotes tumor cell growth

ADAM17 is overexpressed in many human cancers, including lung cancer, while genetic or pharmacological ablation of ADAM17 suppressed tumor proliferation and dissemination partly by promoting EGFR ligand release (McGowan *et al*., 2008; Jiao *et al*., 2018; Saad *et al*., 2019a, 2019b; Ni *et al*., 2020; Schumacher and Rose-John, 2022). Having observed that iTAP expression in recipient host mice affects WT LLC tumor cell growth *in vivo*, we next assessed whether cell-autonomous iTAP expression influences LLC tumor development, this time in WT immunocompetent recipient mice (Fig. 4). To this end, we generated iTAP KO LLC cells (KO LLC) via CRISPR, as well as LLC cells overexpressing iTAP. In pairwise experiments, we confirmed that the levels of ADAM17 maturation were decreased in iTAP KO cells compared to control cells (parental cells expressing Cas9) (Fig. 4A-B) and that iTAP overexpression compared to cells that were transduced with the respective retroviral empty vector enhanced ADAM17 maturation (Fig. 4C,D). Prior to animal studies, we carried out an *in vitro* characterization of these cell lines to test whether the modulation of iTAP levels in LLC cells could impact upon the shedding of the ADAM17 substrates TNF, HB-EGF and Amphiregulin, that could potentially promote differences in cell proliferation and/or migration in *in vivo* experiments. We did not find detectable levels of TNF or HB-EGF (data not shown), but interestingly the level of Amphiregulin shedding by LLC cells was high (Fig.4 E,F). We found that when the cells were stimulated with PMA, to promote ADAM17 activation, iTAP KO cells shed significantly less Amphiregulin (Fig. 4E), whereas the opposite was found in iTAP-overexpressing cells (Fig. 4F). This effect appeared dependent on ADAM17 since the secretion of Amphiregulin in WT cells was significantly decreased by the metalloprotease inhibitor BB-94, but not with the ADAM10-selective inhibitor GI254023X (Fig. 4E,F). Our data establish that iTAP regulates the shedding of endogenous EGFR ligands in cancer cells.

**Figure 4.**
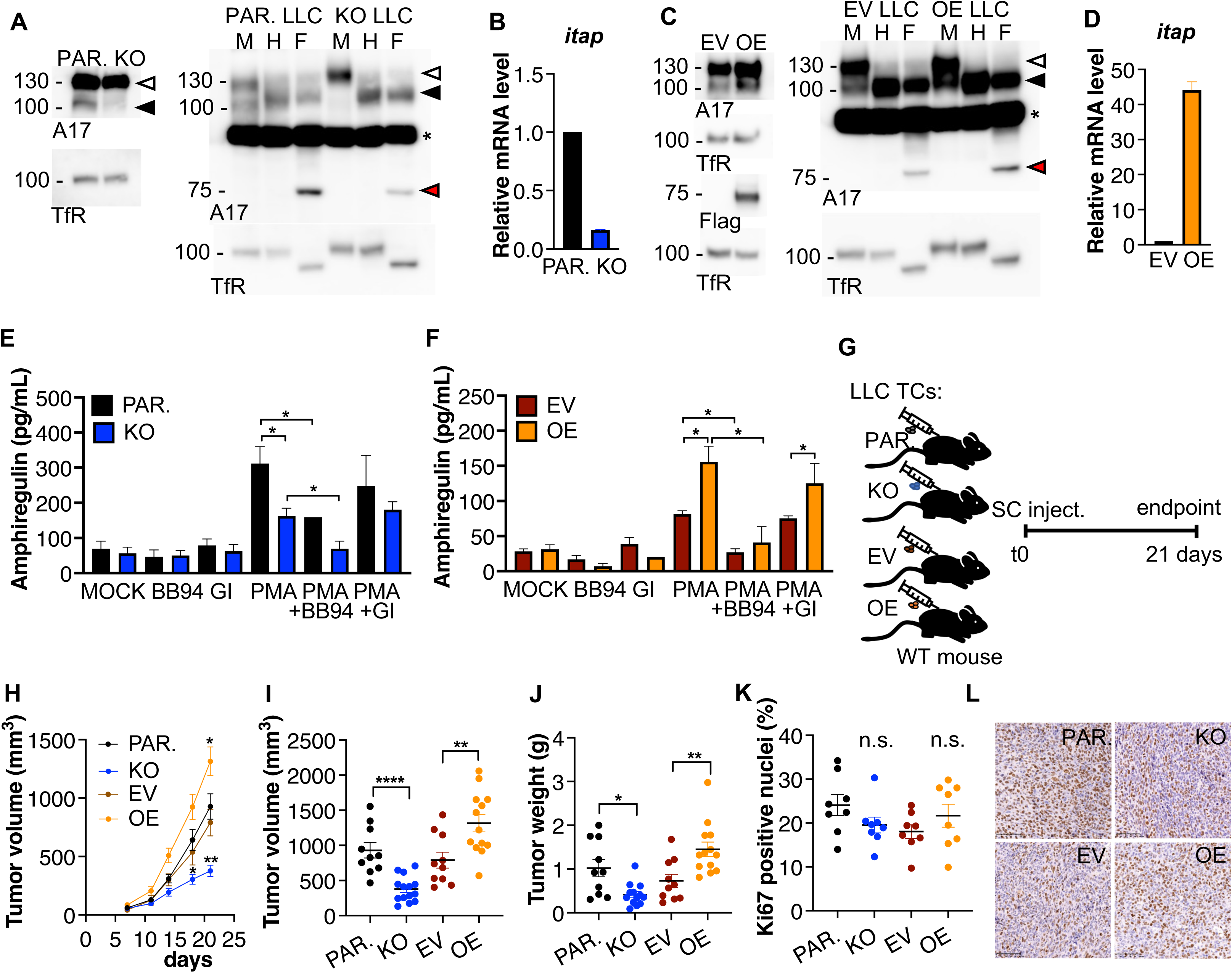
Cell-autonomous expression of iTAP promotes tumor growth. **A, C** Immunoblots showing expression levels of immature versus mature ADAM17 (A17) in parental (PAR.) versus iTAP KO LLC cells (A) or empty vector (EV)- transduced versus iTAP overexpressing (OE) LLC cells (C). Protein samples were deglycosylated with Endo-H (H) and PNGase F (F) versus Mock (M). Immunoblots for the Transferrin receptor (TfR) serve as a loading control. The immature ADAM17 is indicated with white arrowheads; the black arrowhead denotes both glycosylated mature ADAM17 and deglycosylated immature ADAM17 respectively (which have similar electrophoretic mobility), whereas red arrowheads denote the fully deglycosylated, mature, ADAM17 polypeptide. The asterisk indicates an unspecific band. **B, D** Relative mRNA expression levels of iTAP in parental versus iTAP KO LLC cells (B) and empty vector (EV)-transduced control cells versus iTAP-overexpressing (OE) LLC cells (D). Expression was normalized to the levels of a housekeeping control mRNA (*Gapdh*). **E-F** Soluble Amphiregulin levels in the medium of parental versus iTAP KO cells (E) or empty vector (EV) control cells versus iTAP-overexpressing (OE) LLC cells (F). The cells were pre-incubated with DMSO, the metalloprotease inhibitor BB-94 or the ADAM10-specific inhibitor GI254023X (GI) and then treated with DMSO or PMA. n=3. **G** Schematic illustrating the protocol for the subcutaneous (SC) tumor model. **H** Volume of subcutaneous tumors in WT immunocompetent mice receiving iTAP KO LLC cells or iTAP-overexpressing (OE) LLC cells over time compared to their respective controls. **I-J** Volume (I), and tumor weight (J) of the tumors described above **K-L** Level of ki67-positive nuclei (K) and representative images (L) of the tumors described above. n=3 with 3-6 mice per genotype per experiment. For ki67 analysis n=3 with 2-3 mice per genotype per experiment. Results are indicated as mean ± SEM. A scale bar is indicated within the ki67 IHC images. * represents p<0.05, ** represents p<0.01, **** represents p<0.0001.

We next tested the impact of these cells when injected subcutaneously into immunocompetent WT mice (Fig. 4G), finding that the local tumors in the mice that received the iTAP KO cells had a reduced growth rate, leading to smaller tumor volume and mass compared to parental controls (Fig.4 H-J). The opposite was observed in the mice that received the iTAP-overexpressing LLC cells compared to vector controls (Fig.4 H-J). Consistent with these findings, iTAP KO tumors had a tendency towards fewer proliferating cells, as assessed by Ki67 staining; the opposite was found in iTAP-overexpressing tumors (Fig. 4K,L). Despite the modest level of lung metastasis observed in these experiments, the mice that received iTAP KO cells had no lung micrometastases (Fig. S5A,B), while the ones that received the iTAP- overexpressing cells exhibited the opposite trend, presenting more lung micrometastases than controls (Fig. S5C,D).

### Cell-autonomous iTAP expression promotes tumor cell dissemination

As ADAM17 overexpression is associated with higher risk of metastasis and a poorer prognosis in several types of cancer (McGowan *et al*., 2008; Jiao *et al*., 2018; Ni *et al*., 2020), we next determined the cell-autonomous impact of iTAP upon tumor emergence in the lungs following the intravenous injection into WT mice of iTAP KO LLC cells versus their appropriate parental control line (Fig. 5A-E). In separate pairwise experiments, we also compared the injection of empty vector-bearing versus iTAP-overexpressing cells (Fig. 5F-J). In this model, loss of ITAP significantly reduced the number of lung tumors (Fig. 5B), as well as the tumor volume (Fig. 5C) and burden (Fig. 5D) compared to parental WT cells. Consistent with this, the opposite trend was observed when comparing empty vector-transduced cells versus their respective iTAP-overexpressing cells (Fig. 5F-J).

**Figure 5.**
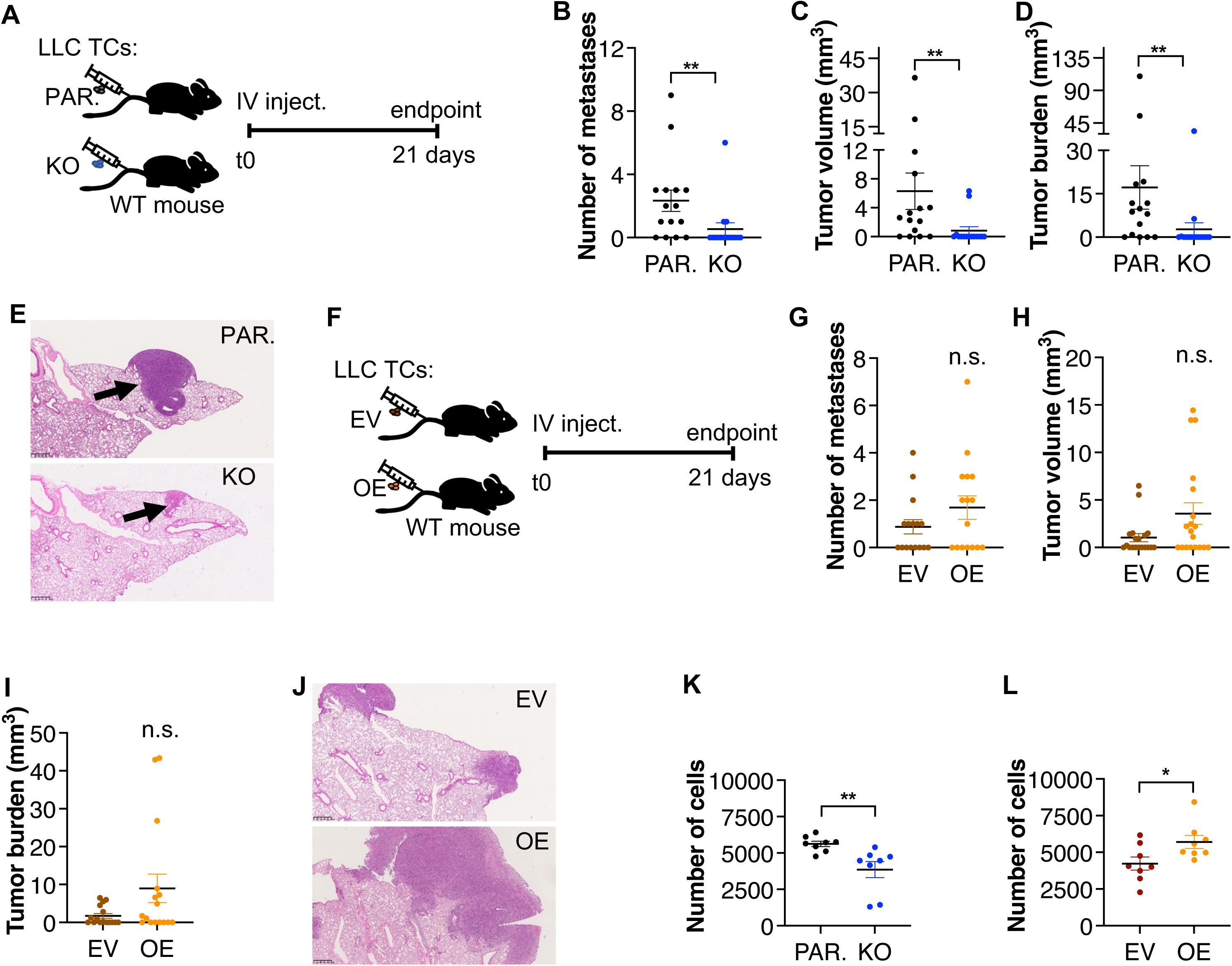
iTAP promotes LLC tumor cell proliferation and metastasis. **A-J** Schematic of the tumor model triggered by *i.v.* injection of parental (PAR.) versus iTAP KO LLC cells into immunocompetent WT hosts (A) or *i.v.* injection of empty vector-transduced versus iTAP-overexpressing (OE) LLC cells into immunocompetent WT hosts (F). The number of lung metastases (B,G), mean metastatic tumor volume (C,H), tumor burden (D,I), and H&E images (E,J) isolated from WT mice following tail vein injection with parental versus iTAP KO (A-E) or empty vector-transduced versus iTAP-overexpressing (OE) (F-J) LLC cells. n=3 with 3-8 mice per group. **K**-L Absolute number of parental versus iTAP KO (K) or empty vector-transduced versus iTAP- overexpressing (OE) LLC cells (L) following 72h of culture with medium supplemented with 0.5% FCS. Results are indicated as mean ± SEM. A scale bar is indicated within the H&E images. * represents p<0.05, ** represents p<0.01.

As our results (Fig. 4 and Fig. 5A-J) indicate that iTAP can promote LLC tumor cell growth and proliferation, we turned to simpler *in vitro* models to explore this effect. We confirmed *in vitro* in pairwise experiments that the iTAP KO LLC cells proliferate more slowly than parental control cells (Fig. 5K), whereas iTAP-overexpressing cells proliferated more rapidly than their equivalent empty vector-transduced cells. (Fig. 5K,L). Taken together with the experiments shown in Figs. 3 and 4, our data establish that iTAP influences tumor growth in two ways: in a cell-autonomous manner that appears to involve control of cellular proliferation, as well as a cell-non-autonomous manner, by influencing the tumor niche.

## Discussion

Studies on iTAP, performed by our laboratory and others revealed that this molecule binds to iRhom proteins and is crucial to stabilize the sheddase complex on the cell surface, avoiding its mis-sorting to and degradation in lysosomes (Künzel *et al*., 2018; Oikonomidi *et al*., 2018). However, the pathophysiological role(s) of iTAP were not fully addressed until now. Our study identifies iTAP as an *in vivo* regulator of ADAM17: in addition to regulating the levels of mature ADAM17 in multiple tissues (Oikonomidi *et al*., 2018) (Fig.1; Fig. S1; Fig. S2), iTAP KO mice exhibit phenotypes associated with ADAM17 shedding defects in several immune cell compartments. This includes reduced shedding of the ADAM17 substrate L-selectin (Marczynska *et al*., 2014) from T lymphocytes and B cells (Fig. 1) and impaired LPS-triggered release of TNF from macrophages *ex vivo* and *in vivo* (Fig. 2) which depends on ADAM17(Bell *et al*., 2007; Horiuchi *et al*., 2007). Equivalent shedding defects have been reported for iRhom2 mutant mice, which also develop normally, but have TNF-associated defects upon inflammatory challenge with LPS (Adrain *et al*., 2012; McIlwain *et al*., 2012; Siggs *et al*., 2012).

Our observation that iTAP KO mice exhibit a normal intestinal epithelium under control conditions but are sensitized to DSS-mediated colitis is consistent with the observation that mice homozygous for a hypomorphic mutation of ADAM17 (ADAM17^ex/ex^) are defective in epithelial repair responses in response to DSS because of a failure to shed EGFR ligands (Chalaris *et al*., 2010). In agreement with this, the shedding of endogenous levels of the key EGFR ligand amphiregulin is defective in iTAP KO LLC cells, and increased in cells overexpressing iTAP (Fig.4).

An obvious question concerns why on one hand iTAP KO mice are ostensibly normal unless challenged with inflammatory insults or cancer models, yet on the other hand they exhibit substantially reduced levels of mature ADAM17 across a range of tissues. This includes tissues in which ADAM17 mutant mice themselves exhibit profound developmental phenotypes, including the heart, skin/keratinocytes (Fig. S1), liver and lung (Oikonomidi *et al*., 2018). One possibility is that the traces of mature ADAM17 that persist in iTAP KO tissues are sufficient to prevent the pronounced embryonic phenotypes exhibited in ADAM17 KO or iRhom DKO models. Indeed, ADAM17^ex/ex^ mice, a hypomorphic mutant that expresses traces of ADAM17, are viable but exhibit eye, skin and heart defects (Chalaris *et al*., 2010), indicating that mouse development is extremely sensitive to threshold levels of ADAM17 activity. Consistent with this notion, ADAM17^ex/ex^ mice are more profoundly sensitized to DSS-induced colitis (Chalaris *et al*., 2010) than appears to be the case in iTAP KO mice (Fig. 2), suggesting that the titre of mature ADAM17 available may be higher in iTAP KO mice than in ADAM17^ex/ex^ mice. Potentially, the residual traces of mature ADAM17 that survive in iTAP KO cells could be the result of redundancy between iTAP and other FERM domain-containing molecules that may bind to and regulate the trafficking of the sheddase complex. Further studies will be required to clarify this. Given that the most pronounced impact of loss of iTAP is in immune cells, it will be interesting to establish whether loss of iTAP has a more pronounced impact on iRhom2-containing sheddase complexes compared to iRhom1-containing sheddase complexes. This could be reconciled, for example, by differential redundancy between iTAP and other FERM domain-containing proteins in tissues that predominantly express iRhom1 versus iRhom2. While a preferential impact of iTAP on iRhom2 biology is consistent with the lack of impact on mouse development and the inflammatory defects associated with iRhom2 KO mice, iTAP binds to iRhom1 (Künzel *et al*., 2018; Oikonomidi *et al*., 2018) and ADAM17 maturation defects have been observed in lysates from the brains of iTAP KO mice (Künzel *et al*., 2018), where iRhom1 is believed to play a more prominent role. Further studies are required to establish the relative impact of loss of iTAP upon iRhom1 versus iRhom2; and to clarify whether additional candidate iRhom-interacting FERM domain proteins fulfil redundant roles in some tissues.

An alternative explanation for the mild phenotypes observed in iTAP KO mice could reflect the compartment within the cell where loss of iTAP exerts its effects on the sheddase complex. Whereas KO (or DKO) of iRhom proteins results in a trafficking block early in the secretory pathway that prevents the generation of mature ADAM17 (Adrain and Freeman, 2012; Christova *et al*., 2013; McIlwain *et al*., 2012; Li *et al*., 2015), depletion of iTAP triggers the precocious mis-sorting of the sheddase complex to lysosomes, wherein iRhom and ADAM17 are degraded (Künzel *et al*., 2018; Oikonomidi *et al*., 2018). As iTAP appears to influence sheddase complex trafficking within the endocytic or endolysosomal pathway, iTAP’s influence occurs after mature ADAM17 has been generated. Speculatively, this could allow small steady state levels of ADAM17 substrate shedding to occur on or near the cell surface prior to the sheddase complex being mis-sorted to and degraded in lysosomes. This transient exposure of mature ADAM17 to its substrates could prevent the catastrophic loss of ADAM17 function phenotypes. Future studies will be required to define the precise trafficking route taken by the sheddase complex in normal cells and the defective itinerary incurred in iTAP KOs.

Our study also establishes that iTAP expression can influence multiple aspects of tumor growth and dissemination: it conditions the host microenvironment to promote tumor growth and the formation of micrometastases (Fig. 3), while also influencing cell-autonomous tumor growth and dissemination (Fig. 4, Fig. 5). These observations are potentially consistent with ADAM17-associated defects in the cleavage of pro-inflammatory factors (e.g. TNF, IL6R (Schumacher and Rose-John, 2022)) that can recruit inflammatory cells to the tumor microenvironment to drive proliferation of tumor cells, or the release of growth factors such as Amphiregulin (Fig. 4E,F) to promote tumor cell growth Interestingly, mutations or copy number variations within the *FRMD8* gene have been reported in clones isolated from patients suffering from myeloid malignancies. This includes the pre-leukemic condition myelodysplastic syndrome, where *FRMD8* mutations have been found in patient clones following chemotherapy(da Silva-Coelho *et al*., 2017). Moreover, amplifications in *FRMD8* copy number have been observed in pediatric acute myeloid leukemia patients, particularly within the context of primary chemotherapy resistance (McNeer *et al*., 2019). Finally, in adult AML *FRMD8* overexpression is associated with a poorer prognosis (Bou Samra *et al*., 2012). It will be interesting to determine, in future studies, whether regulation of the sheddase complex underpins the association between iTAP and poor cancer prognosis, or whether iTAP regulates additional pathways within the context of myeloid malignancies.

One of the major challenges in the potential therapeutic modulation of ADAM17 activity concerns the poor preclinical performance (Murumkar *et al*., 2020; Calligaris *et al*., 2021) of numerous ADAM17-specific drugs which have proved to be toxic, in part because of cross-reactivity with related metalloproteases. Indeed, given the crucial biological roles fulfilled by ADAM17 in multiple organs during development, it is arguably desirable to achieve partial–rather than complete–therapeutic ADAM17 inhibition, or indeed, to achieve tissue-specific blockade of ADAM17 activity. With this in mind, the targeting of iTAP may be an appealing potential strategy to target ADAM17 activity in specific compartments during chronic inflammatory diseases or cancer, while avoiding collateral impact on the vital functions of ADAM17 in normal tissues.

## Experimental procedures

### Experimental animals

iTAP KO mice were generated as previously described on a C57BL/6J background (Oikonomidi *et al*., 2018). Mice were maintained in a SPF facility on a 12-hour light/dark cycle, at standard sub-thermoneutral conditions of 20-24°C and an average of 50% humidity, in ventilated cages with corn cob as bedding. We co-housed C57BL/6J WT and ITAP KO animals together to homogenize differences in microbiota and other environmental conditions. Food intake and body weight was recorded weekly from the age of 5 weeks and for 9-10 weeks.

### Mouse tumor model

Two cancer models were used, based on the inoculation of LLC (Lewis lung carcinoma, ATCC# CLR1642) cells. To assess topical growth 0.5 million LLC cells in PBS were inoculated subcutaneously (100 μL) into the flank of WT versus iTAP KO mice. Tumor dimensions were measured with a caliper one week after inoculation, and afterwards twice a week until 21 days after, when the mice were euthanized and the tumor and lungs were collected. Tumor weight, number, volume and burden was evaluated, as was the presence of metastasis in the lungs as previously described(Mendonça *et al*., 2019). To assess LLC cell migration to the lung, in a second model, 0,5 million LLC cells in PBS were inoculated intravenously into the tail vein (200 μL) and after 21 days the mice were euthanized, after which the lungs were collected and their tumors counted and measured. For both models, tumor volume was calculated using the formula V= 0,52 x a x b^2^, where a and b equal the longer and shorter length of the tumor, respectively.

### Acute colitis model

To induce intestinal inflammation, 8-12 weeks old iTAP KO and WT mice were given 3 % Dextran sulfate sodium salt (DSS) in the drinking water for a total of 8 days. The DSS solution was replenished every 2 days. Body weight was measured daily. At day 8, water was given to the mice and at day 10 the mice were euthanized and the large intestine was collected, and its length was measured as described (Wirtz *et al*., 2017).

### Sepsis model

Lipopolysaccharide (LPS) was injected intraperitoneally into 8-12 weeks old iTAP KO and WT mice at a dosage of 37.5 mg/Kg in PBS. Serum was collected via cardiac puncture from animals that only received PBS (0), and from mice administrated with LPS at 3h and 6h post-LPS injection. TNF and IL-6 levels were measured from serum using specific ELISA kits (88-7324-22, 88-7064-88, respectively, eBioscience).

### Western blotting

Cells isolated from iTAP KO and WT mice were washed in PBS and lysed for 10 minutes on ice in TX-100 lysis buffer (1% Triton X-100, 150 mM NaCl, 50 mM Tris-HCl, pH 7.4) containing protease inhibitors and 0.1 mM of the metalloprotease inhibitor 1,10-phenanthroline. 1,10-Phenathroline was added because, without it ADAM17 autocatalytically cleaves off its cytoplasmic tail, resulting in the loss of the epitope detected by the anti-ADAM17 antibody (Ab39162, Abcam) (Adrain *et al*., 2012). Tissues from the mice described above were lysed in a modified RIPA buffer (1% Triton X-100, 150 mM NaCl, 50 mM Tris-HCl, pH 7.4, 1 mM EDTA, 1 % Sodium Deoxycholate, 0.1 % SDS) containing protease inhibitors and 0.1mM of 1,10- phenanthroline using a TissueLyser II (QIAGEN). The samples were then quantified, normalized, and to improve the detection of ADAM17, glycoproteins were captured using 40 μl of Concanavalin A (ConA) Agarose. Beads were washed twice in the same buffer supplemented with 1 mM EDTA, 1 mM MnCl2, 1 mM CaCl2 and eluted by heating for 15 min at 65 °C in sample buffer supplemented with 15 % sucrose. After centrifugation at 1000 g for 2 min the supernatants where collected. In some cases, the denatured lysates were digested for 2 hr at 37 °C with deglycosylating enzymes Endoglycosidase-H (Endo-H), which removes high mannose N-linked glycans added in the ER, but not complex N-linked glycans found in the later secretory pathway, and with PNGase F, which deglycosylates both. Then, the samples were denatured for 5 min at 65 °C. The lysates were fractionated by SDS-PAGE and transferred onto PVDF membranes. After blocking with 5 % milk in TBS-T for 30 min, the membranes were cut and incubated overnight with the primary antibodies rabbit anti-ADAM17 antibody (1:1000, Ab39162, Abcam), mouse anti-Flag HRP (1:1000; A8592; Sigma), rat anti-tubulin (1:1000 Clone YL1/2, IGC antibody facility), mouse anti-Transferrin receptor (1:1000, 13-6800, Life Technologies). The following day, the membranes were washed and incubated with the secondary antibody anti-rabbit HRP (1:5000; 1677074P2, Cell Signaling Technology), anti-mouse HRP (1:5000; 1677076P2, Cell Signaling Technology) or anti-rat HRP (IGC Antibody Facility). After washing, protein bands were detected using ECL.

### Histopathological analysis

Histopathological analysis was performed by a veterinary pathologist in a blinded manner. Sections were examined under a Leica DMLB2 microscope. Collected samples were formalin-fixed, paraffin embedded, sectioned (3 μm sections) and stained with H&E (Sigma). LLC tumor samples and dorsal skin were embedded in 3.5- 4 % agar. Each specimen was sectioned into a minimum of 8 slices of equal thickness. Depending on the size of the sample, they were sectioned into 1.3 mm or 2.6 mm slices. Whole slide images were acquired with Hammatsu Nanozoomer slide scanner. Measurements were performed with the Visiopharm stereology software. Volume was estimated according to the Cavalieri principle, in which a grid of equally spaced points is overlayed onto systematic fields of view, automatically generated by the software. Each point has an associated area. Every time a point hits the tissue of interest, it is counted. The volume of the tissue of interest equals the number of points (Qi) multiplied by the area per point (a/p), multiplied by the section thickness. V = Qi x a/p x T. Lungs were exhaustively sectioned and every 30^th^ section a slice was chosen, resulting in a total of around 20 sections per lung, separated by 90 μm. A grid of 5x5 points was used for the tumors and of 3x3 points was used to evaluate the lungs. In the case of the LLC allograft model, every time a point hit necrosis or superficial ulceration, it was counted separately. The left-hand top corner point was used for tumor estimation (the area of this point is 25 times larger than the rest of the points). The measurements were done across 20-30 % of the section’s region of interest in fields of view with 10X magnification for the tumors, and 5X magnification for the lungs. The volume of lung micrometastases was estimated with a grid of 15x15 points, across 50 % of the section’s region of interest in fields of view with 20X magnification.

The large intestines from the colitis model were scored as described (Seamons *et al*., 2013). Specifically, the proximal, mid and distal colon was scored by the level of mucosal loss (0 none, 1 mild <5 %, 2 moderate 6-30 %, 3 marked 31-60 %, 4 severe >61 %), hyperplasia (0 none, 1 mild, 2 moderate, 3 severe with aberrant crypts, 4 severe with herniation), inflammation (0 none, 1 mild mucosa only, 2 moderate mucosa and submucosa, 3 severe with abscesses, erosion, 4 severe with obliteration of architecture and ulceration or transmural), extent of intestine affected and of the intestine affected by the most severe score (0 none, 1 <5 %, 2 6-30 %, 3 31-60 %, 4 >60 %). The IBD total score resulted of the sum of the scores for the proximal, mid and distal colon.

The WT or iTAP KO dorsal skin was sectioned at 250 μm, resulting in a total of 5 to 8 slabs per skin. The slabs were then paraffin-embedded and photographed for volume estimation by point counting as described above. Eight paraffin sections (5 μm) were sampled from the upper-cut surface of each slab and stained with H&E. Estimations of follicle number were performed using the dissector principle, where the distance between the reference and the look-up section must represent 30 % of the object’s height. In this case, optimal distance was found to be in range of 30 to 40 μm. If a particle is seen in only one of the sections, it is counted. The number of differences between the upper and lower sections is given by 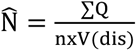, where V(dis) is the reference volume and n is the number of sections. Follicle number was obtained by multiplying the numerical density by the total skin volume. Estimates of skin and epidermal volume were obtained according to the Cavalieri principle (Gundersen and Jensen, 1987). V = T x a/p x ∑Pi, where T is the thickness of the slabs, Pi is the number of points hitting the structures of interest and (a/p) is the area associated with each point. After paraffin embedding and due to expected tissue shrinkage, thickness was remeasured. Epidermal fraction was estimated dividing the points hitting the epidermis by the points hitting the skin. V = Pi(Epidermis)/Pi(Skin). To estimate surface area, a cycloidal test grid was superposed on the sections. A reference volume was obtained by the Cavalieri principle. Surface density was then estimated by counting the number of intersections (I) between the cycloids and the epidermal/dermal interface. 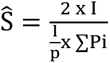 where I is the number of intersections counted, Pi is the number of points counted and l/p is the test line length per point at the level of the tissue. Surface area was obtained by multiplying the surface density by the total skin volume. To facilitate interpretation, all previous measurements were converted into structure of interest per mm³. The stereological measurements were done with the software STEPanizer.

### Immunohistochemistry

Tumors samples were deparaffinized and hydrated. Antigen retrieval was performed with Citrate buffer 50 min at 99 °C in a water bath. Cooled samples were exposed to 3% hydrogen peroxidase solution for 10 min, washed with PBS-Tween 20 and exposed to M.O.M. Mouse Ig Blocking Reagent (MKB-2213-1, Vector Laboratories) for 1 h. After 2 washes, the samples were incubated with the primary antibody mouse anti–Ki67 (1:200, 550609, BD Pharmigen) for 1 h. After 2 washes, the samples were incubated with the respective M.O.M. ImmPRESS Anti-Mouse IgG Reagent (MPX- 2402-15, Vector Laboratories) for 10 min. After 2 washes, the samples were incubated with DAB (K3468, Dako) for 8 min, washed, and then incubated with Mayer’s haematoxylin (HMS 16, Sigma) for 4 min. Then, the samples were washed, dehydrated, cleared and mounted.

Computer-assisted semi-quantitative analysis was used to evaluate Ki67 ratio using Image J plugin Immunoratio. The data was expressed as the % of Ki67 positive nuclei in 5 different pictures taken at 20x. A Macro was used to measure DAB, that can be provided upon request.

### Generation of iTAP KO and overexpressing (OE) LLC cell lines

Lewis lung carcinoma (LLC) cells were cultured in DMEM high glucose (Biowest), supplemented with 10% FCS (Biowest) and 1% Pen/Strep (Sigma).

To generate an iTAP KO LLC cell line, sgRNAs targeting coding exon 3 (5’CCAGACCTTACCCAAGAGAG 3’, 5’ CGATGTCCTGGTGTACCTGG 3’) and exon 4 (5’ CATGGCGACATCATCATCGG 3)’, were cloned into plasmid px330 (Zhang lab, 42230, Addgene). HEK 293FT cells (6 x 10^6^) were transfected with pMD-VSVG envelope plasmid, psPAX2 helper plasmid, and pLentiCas9-Blast plasmid (52962, Addgene) or pLentiGuide-puro (52963, Addgene) plus the 3 iTAP sgRNA sequences inserted into CRISPR plasmid px330 using Fugene HD-based lipofection. After 48h the virus was collected. Ultracentrifuged, 150x concentrated lentivirus LentiCas9-Blast (20 μl) was added to 180,000 LLC cells supplemented with 8 μg/mL polybrene. After selection with blasticidin (16 μg/mL) the cells were left for 1 week to allow Cas9 expression. Then, lentivirus LentiGuide-puro plus the three iTAP sgRNAs were added (2 mL, in two rounds with a 24 h interval) to 180,000 LLC cells expressing lenti-Cas9 supplemented with polybrene. The cells where then selected with puromycin (6 μg/mL) and plated in a limiting dilution in 96 well plates to isolate single clones. The clones were screened by qPCR to confirm depletion of iTAP mRNA levels. Three clones with the strongest iTAP deletion were combined into a pool and the parental Cas9- expressing cells were used as controls.

To generate a LLC cell line overexpressing iTAP, HEK 293ET cells (6 x 10^6^) were transfected with the plasmids pCL-Eco packaging plasmid (Naviaux *et al*., 1996) plus pM6P.BLAST empty vector (kind gift of F. Randow, Cambridge, UK), or pM6P containing the mouse iTAP WT cDNA fused to a Flag tag using Fugene HD-based lipofection. After 48h the retroviruses were collected and added (2 mL, twice sequentially, with a 24 h interval) to 180,000 LLC cells supplemented with polybrene 8 μg/mL. The cells where then selected with puromycin (6 μg/mL). The impact upon ADAM17 maturation and FRMD8 mRNA levels were assessed as by western blotting and qPCR, respectively. These cell lines were then used in mouse experiments.

LLC iTAP KO and OE versus their respective controls (PAR-parental and EV-empty vector) were plated in a 24 well plate at a density of 700,000. After 4 h the cells were starved in serum free medium overnight and then incubated for 1h with the ADAM17 inhibitor Batimastat (BB-94) at 10 μM (196440, Calbiochem), with the selective ADAM10 inhibitor GI254023X (GI) at 1 μM (SML0789, Sigma) or Dimethyl sulfoxide (DMSO) (67-68-5, Sigma). Then the cells were stimulated with Phorbol-12-myristate- 13-acetate (PMA) at 1 μM (P1585, Sigma) or DMSO for 1h. The medium as collected and after clearance, where used to evaluate the levels of TNF, HB-EGF and Amphiregulin using specific ELISA kits (88-7324-22 eBioscience, DY8239-05, R&D Systems, DY989, R&D Systems, respectively).

### Isolation and differentiation of primary cells in vitro

Bone Marrow Derived Macrophages and Dendritic cells (BMDM and BMDC, respectively) were isolated from 8-10 week old iTAP KO and WT mice tibias and femurs and BMDM were differentiated as previously described(Adrain *et al*., 2012) in RPMI 1640 with Glutamax (Biowest), supplemented with 10% FCS (Biowest), 1% Pen/Strep and gentamicin sulphate (10 μg/mL) (Sigma), 20% of L929 cell (CCL-1, ATCC) conditioned medium, and 50 μM 2-Mercaptoethanol (Sigma). At day 7 of differentiation 0,5 million cells were plated in a 24 well plate. At day 8 cells were stimulated with LPS at 0.25, 0.5 and 1 μg/mL and the supernatant was collected after 2h and 6h. TNF and IL-6 was measured from clarified supernatants using specific ELISA kits (88-7324-22, 88-7064-88, respectively, eBioscience). BMDCs were differentiated for 8 days in RPMI 1640 with Glutamax (Biowest), supplemented with 10 % FCS (Biowest), 1% Pen/Strep (Sigma), 20 ng/mL GM-CSF, and 50 μM 2- Mercaptoethanol (Sigma). Keratinocytes were isolated from 1-2 days old iTAP KO and WT neo-natal epidermis following the protocol of CELLmTEC. Briefly, the isolated skin was incubated overnight at 4 °C in a tube containing 10 mL of 1x Dispase (Roche) and 2x Pen/Strep (Sigma) and 2x Amphotericin B (Life Tech). The next day, the skin was transferred to CnT-07 medium (CELLmTEC) and the dermis was separated. Accutase (CnT-Accutase-100, CELLmTEC) was then added to the epidermis for a period of 20- 30 min at room temperature. CnT-07 medium was then added and the epidermis was separated into single cells. After washing and resuspension, the cells were cultured in CnT-07 medium supplemented with IsoBoost (CnT-ISO-50, CELLmTEC) for 3 days.

### Flow Cytometry

Spleens and mesenteric, inguinal, cervical and axillar lymph nodes were mashed through a 70 μm cell strainer. The digested cells were centrifuged (5 min, 4 °C, 2000 rpm), and red blood lysis buffer was added (0,88 % NH4Cl). The cells were washed and filtered through a 70 μm strainer. Three million splenocytes were plated per condition in RPMI medium (Biowest) supplemented with 2% FCS (Biowest). Samples were incubated for 1 h with the ADAM17 inhibitor Batimastat (BB-94) at 10 μM (196440, Calbiochem), with the selective ADAM10 inhibitor GI254023X (GI) at 1 μM (SML0789, Sigma) or Dimethyl sulfoxide (DMSO) (67-68-5, Sigma). Afterwards the cells were stimulated with Phorbol-12-myristate-13-acetate (PMA) at 1 μM (P1585, Sigma) or LPS (0.25 μg/mL) or DMSO for 2h. Samples were centrifuged and resuspended in PBS supplemented with 2% FCS and incubated in Fc Block, clone 2.4G2 (1:100, produced in-house) for 15 min. After washing, staining was performed by incubation for 30 min with anti-mouse CD45.2-PE, clone104.2, B220-PE-Cy7, clone RA3-6B2, L-selectin-7AAD, clone MEL-14, CD4-FITC, clone GK1.5, CD8- Pacific Blue, clone YTS169.4, and CD3-A647, clone 500A2 (all at 1:100; produced in-house) and with the Zombie Aqua fixable dye (423101, Biolegend).

For B cell sorting, the total amount of splenocytes and lymphocytes collected were washed and incubated in Fc Block for 15 min. Then the samples were washed and incubated for 30 min with anti-mouse CD45.2-PE, clone104.2, CD19-PE-Cy7, clone 6D5, CD3-A647, clone 500A2 (all at 1:100; produced in-house) and with the Zombie Aqua fixable dye (423101, Biolegend).

The purity of DCs was assessed by staining for anti-mouse CD45.2-A647, clone104.2, CD11b-FITC, clone M1/70 and CD11c-PE, clone HL3 (all at 1:100; produced in-house).

LLC iTAP KO and OE versus their respective controls (PAR. and EV) were plated in 6 wells at a density of 0.5 million cells. After 4 h the cells were starved in serum free medium overnight and afterwards maintained in medium containing 0,5 % FCS for 48h. Cells were trypsinized, washed, and then stained with DAPI (0.2 μg/mL; Invitrogen) to count the absolute number per well by adding Precision Count Beads™ (BioLegend) to the samples. All the samples were analyzed using a Fortessa X20 with FlowJo software, version 10.2.

### Quantitative transcriptional analysis

LLC tumor samples and lungs from iTAP KO and WT mice were snap frozen in liquid nitrogen and kept at -80 °C until RNA extraction (NZYTech). The same was applied to LLC cells. First-strand cDNA was synthesized from total RNA using the SuperScript® III First-Strand Synthesis SuperMix. Real-time PCR analysis was performed using the comparative CT method (Schmittgen and Livak, 2008). Gene expression was normalized to *Gapdh* mRNA levels. Primer sequences are listed in Table 1.

### Statistics

To compare single measurements between control and test groups, Mann-Whitney-Wilcoxon test were used. For repeated measurements, the two-way ANOVA was used. The statistical analysis was performed using GraphPad Prism, version 6. Results are presented as average ± SEM. *P*-values <0.05 were represented as (*), <0.01 as (**), <0.001 as (***).

### Study approval

Animal procedures were approved by the national regulatory agency (DGAV – Direção Geral de Alimentação e Veterinária) and by the Ethics Committee of Instituto Gulbenkian de Ciência and the Institutional Animal Care, and were carried out in accordance with the Portuguese (Decreto-Lei no. 113/2013) and European (directive 2010/63/EU) legislation related to housing, husbandry, and animal welfare.

## Data availability

Data is available within the manuscript.

## Supporting information

This article contains supporting information,

## Acknowledgements

The authors thank the Ethics Committee, Animal Facility, Histopathology and Flow cytometry units and the Antibody Service of the Instituto Gulbenkian de Ciência. We thank Moises Mallo and Ana Nóvoa for advice and help in the generation of iTAP mutant mice. We thank Cristina Branco for reagents. We thank Stefan Rose John and Cristina Branco for advice concerning the LLC cell model and helpful discussions.

## Funding and additional information

We acknowledge the support of Fundação Calouste Gulbenkian; Queen’s University Belfast; Worldwide Cancer Research (14–1289); a Marie Curie Career Integration Grant (project no. 618769); Fundação para a Ciência e Tecnologica (FCT) grants, SFRH/BCC/52507/2014, PTDC/BEX-BCM/3015/2014 and LISBOA-01–0145-FEDER-031330) and funding from ‘La Caixa’ Foundation under the agreement <LCF/PR/HR17/52150018>.This work was developed with the support of the research infrastructure Congento, project LISBOA-01–0145-FEDER-022170, co-financed by Lisboa Regional Operational Programme (Lisboa 2020), under the Portugal 2020 Partnership Agreement, through the European Regional Development Fund (ERDF), and Foundation for Science and Technology (Portugal).

---

**Supplementary Figure 1.**
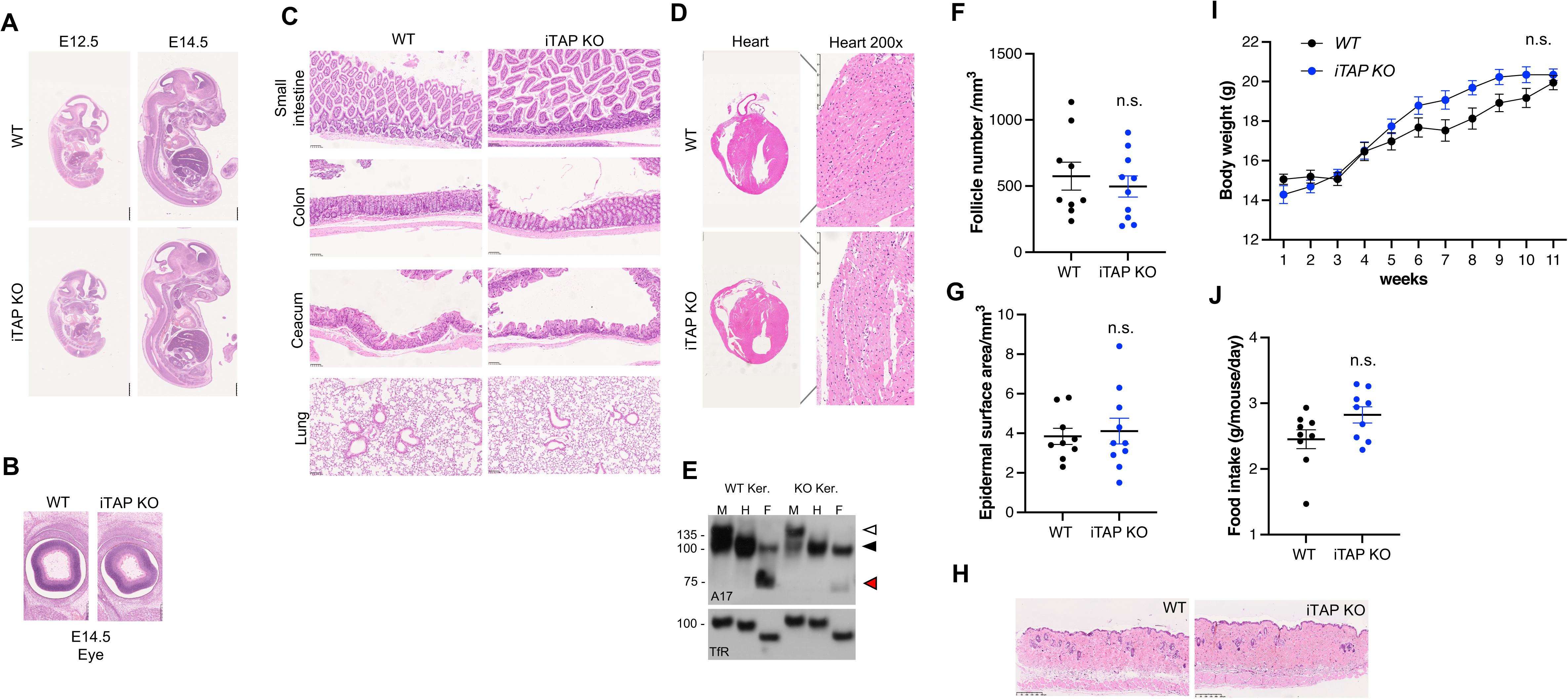
iTAP KO mice develop normally with no major physiological defects. **A-B** H&E staining of WT vs iTAP KO mouse embryos at E12.5 and E14.5 (A) and of the eye at E14.5 (B). **C, D, H** H&E staining of tissues isolated from the small intestine, colon, caecum, lung (C), heart (D), and dorsal skin (H) from WT vs iTAP KO mice. **E** Protein expression of immature and mature ADAM17 in keratinocytes isolated from WT versus iTAP KO mice. Protein samples were either mock-treated (M) or deglycosylated with Endo-H (H) and PNGase F (F). Immunoblots for the Transferrin receptor (TfR) serve as a loading control. n=2 Immature ADAM17 is indicated with white arrowheads; the black arrowhead denotes both glycosylated mature ADAM17 and deglycosylated immature ADAM17 respectively (which have similar electrophoretic mobility), whereas red arrowheads denote the fully deglycosylated, mature, ADAM17 polypeptide. **F-G** Dorsal hair follicle density (F), dorsal epidermis thickness (G). **I, J** Body weight increase over time (I) and food consumption per day (J) of the animals described above. n=6 per genotype.

**Supplementary Figure 2.**
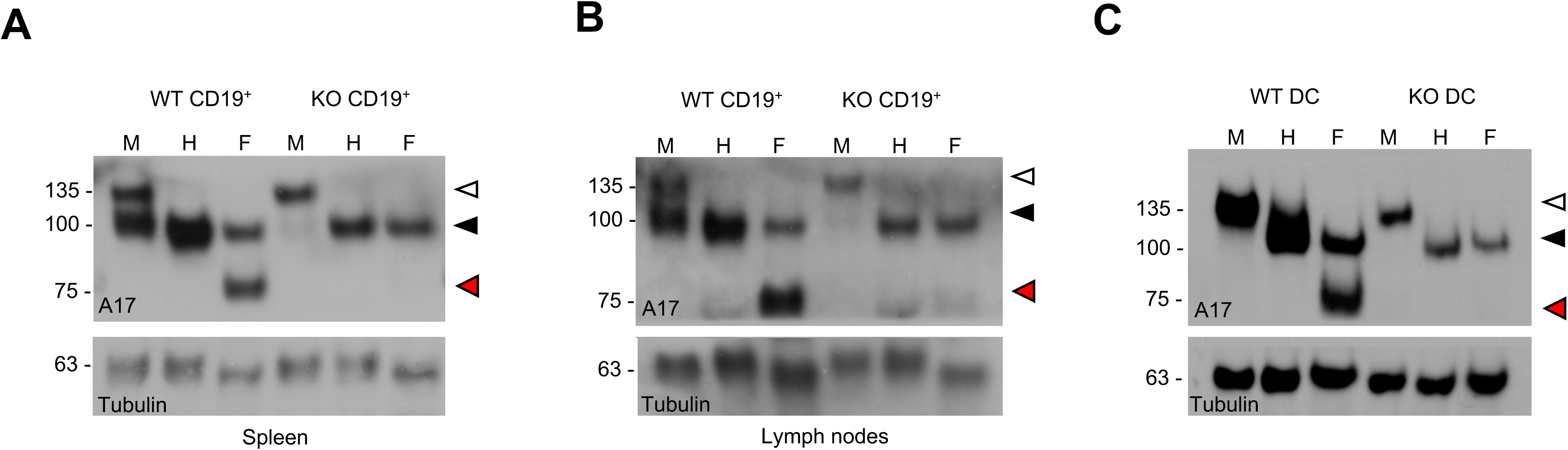
iTAP KO mice have defects in ADAM17 maturation in immune cells. **A-B** Protein expression of immature and mature ADAM17 in B cells (CD45.2^+^, CD19^+^) from spleen (A), and from lymph nodes (B) isolated from iTAP KO mice versus controls and also from dendritic cells (DC) (C) differentiated *in vitro* from bone marrow progenitors. Protein samples were either mock-treated (M) or deglycosylated with Endo-H (H) and PNGase F (F). Immunoblots for Tubulin serve as a loading control. n=2. Immature ADAM17 is indicated with white arrowheads; the black arrowhead denotes both glycosylated mature ADAM17 and deglycosylated immature ADAM17 respectively (which have similar electrophoretic mobility), whereas red arrowheads denote the fully deglycosylated, mature, ADAM17 polypeptide.

**Supplementary Figure 3.**
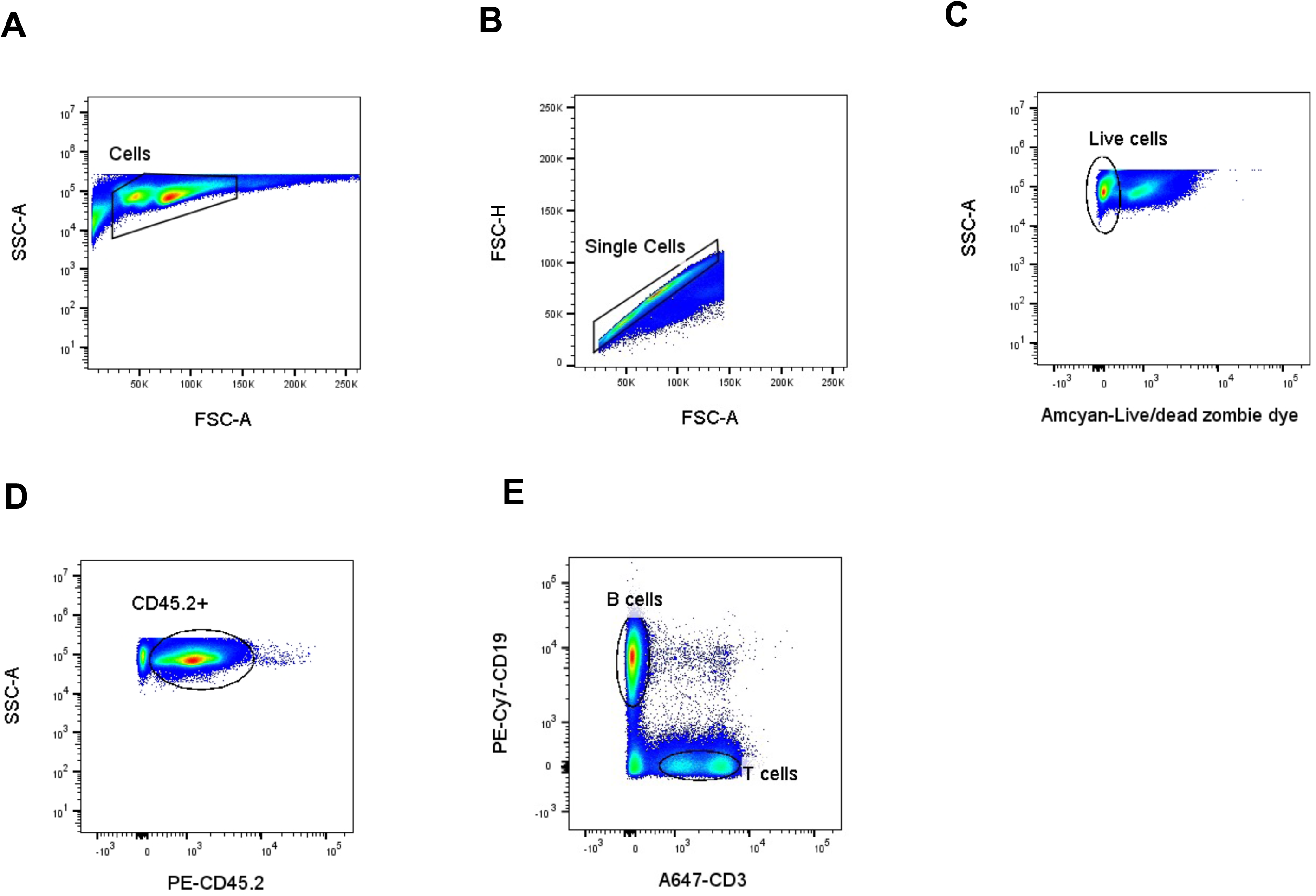
Gating strategy used in the FACS analysis to sort B cells A-E. Cells were first gated for size (SSC-A *vs* FSC-A) (A) and then on singularity (FSC-H *vs* FSC-A) (B). This was followed by Amcvan-Live/dead zombie dye exclusion to gate for live cells (SSC-A *vs* Live/dead dye) (C). Live cells were gated on expression of CD45.2 to select the myeloid cells (SSC-A *vs* PE-CD45.2) (D). Finally, myeloid cells were gated for expression of CD19 and CD3 to identify B and T cells, respectively (PE-Cy7-CD19 *vs* A647-CD3) (E) to isolate the B cell population.

**Supplementary Figure 4.**
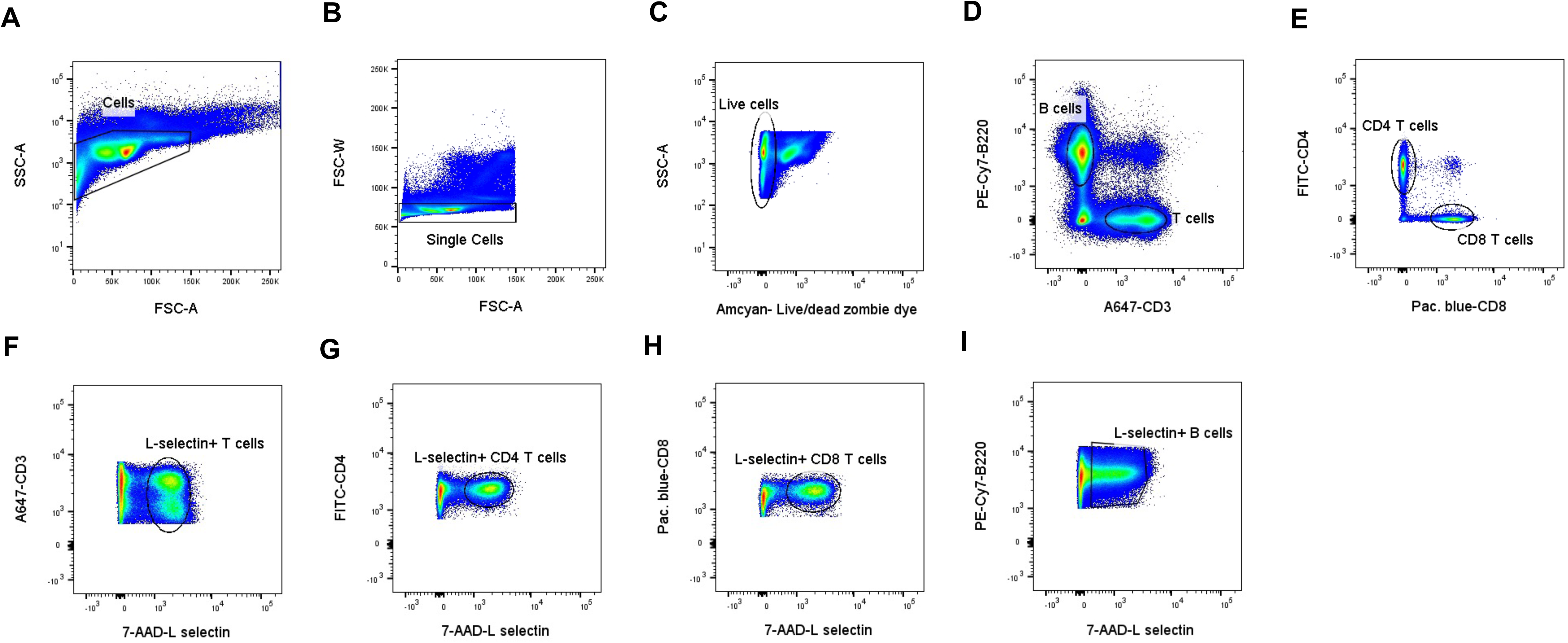
Gating strategy used in the FACS analysis of L-selectin expressing B, and T total and subpopulations cells A-E. Cells were first gated on size (SSC-A *vs* FSC-A) (A) and then on singularity (FSC-W *vs* FSC-A) (B). This was followed by Amcvan-Live/dead zombie dye exclusion to gate for live cells (SSC-A *vs* Live/dead dye) (C). Live cells were gated for expression of B220 and CD3 to identify B and T cells, respectively (PE-Cy7-B220 *vs* A647-CD3) (D). T cells were then gated for expression of CD4 and CD8 to identify helper and cytotoxic T cells, respectively. (FITC-CD4 *vs* Pacific blue-CD8) (E). **F-I** T (F), CD4^+^ T (G) and CD8^+^ T (H) and B (I) cells were then gated for expression of L-selectin at the cell surface (7-AAD-L selectin).

**Supplementary Figure 5.**
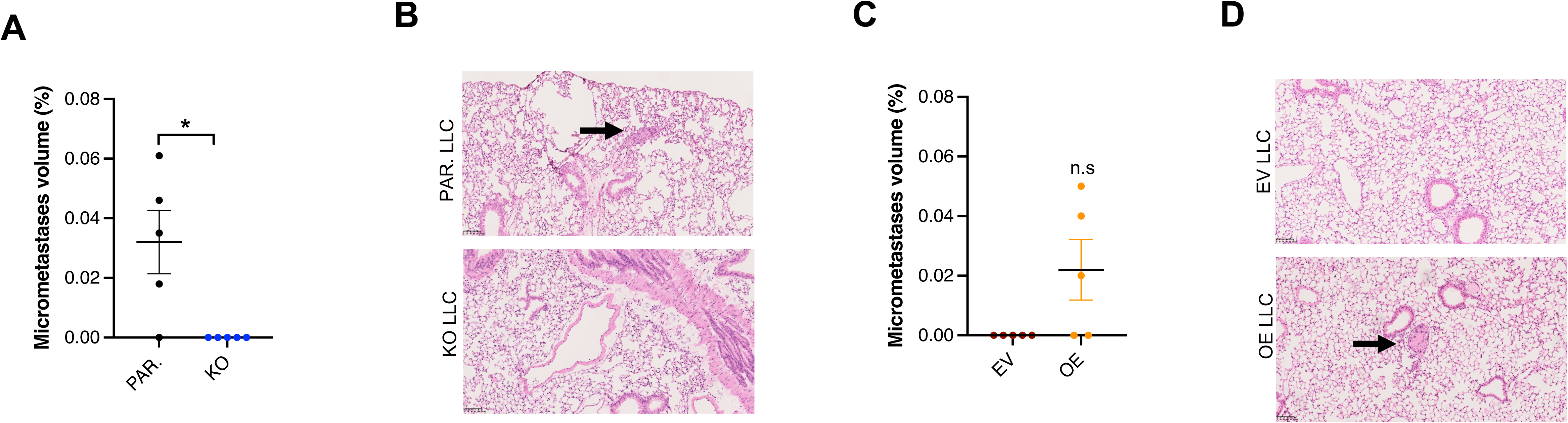
Detection of lung micrometastases in WT mice following injection of iTAP deficient or overexpressing LLC cells A-C. Percentage of micrometastases (A,C) and representative H&E images (B,D) of the lungs of WT mice following subcutaneously injection with parental (PAR.) versus iTAP KO LLC cells (A-B) or empty vector-transduced (EV) versus iTAP- overexpressing (OE) LLC cells (C-D). n=2 with 2-3 mice per group. Results are indicated as mean ± SEM. A scale bar is indicated within the H&E images. * represents p<0.05.

